# Genomic Surveillance of Methicillin-Resistant *Staphylococcus aureus* in the Philippines from 2013-2014

**DOI:** 10.1101/2020.03.19.998401

**Authors:** Melissa L. Masim, Silvia Argimón, Holly O. Espiritu, Mariane A. Magbanua, Marietta L. Lagrada, Agnettah M. Olorosa, Victoria Cohen, June M. Gayeta, Benjamin Jeffrey, Khalil Abudahab, Charmian M. Hufano, Sonia B. Sia, Matthew T.G. Holden, John Stelling, David M. Aanensen, Celia C. Carlos, on behalf of the Philippines Antimicrobial Resistance Surveillance Program

## Abstract

Methicillin Resistant *Staphylococcus aureus* (MRSA) remains one of the leading causes of both nosocomial and community infections worldwide. In the Philippines, MRSA rates have remained above 50% since 2010, but resistance to other antibiotics, including vancomycin, is low. The MRSA burden can be partially attributed to pathogen-specific characteristics of the circulating clones, but little was known about the *S. aureus* circulating clones in the Philippines.

We sequenced the whole genomes of 116 *S. aureus* isolates collected in 2013-2014 by the Antimicrobial Resistance Surveillance Program. The multi-locus sequence type, *spa* type, SCC-*mec* type, presence of antimicrobial resistance (AMR) determinants and virulence genes, and relatedness between the isolates were all derived from the sequence data. The concordance between phenotypic and genotypic resistance was also determined.

The MRSA population in the Philippines was composed of a limited number of genetic clones, including several international epidemic clones, such as CC30-spa-t019-SCC*mec*-IV-PVL+, CC5-SCC*mec*-typeIV, and ST239-spa-t030-SCCmec-typeIII. The CC30 genomes were related to the South West Pacific clone, but formed a distinct and diverse lineage, with evidence of global dissemination. We showed the independent acquisition of resistance to sulfamethoxazole/trimethoprim across different locations and genetic clones, but mostly in pediatric patients with invasive infections. The concordance between phenotypic and genotypic resistance was 99.68% overall for 8 antibiotics in 7 classes.

We produced the first comprehensive genomic survey of *S. aureus* in the Philippines, which bridges the gap in genomic data from the Western Pacific region and will constitute the genetic background to contextualize ongoing prospective surveillance.

## Introduction

Methicillin Resistant *Staphylococcus aureus* (MRSA) remains one of the leading causes of both nosocomial and community infections worldwide. ^1^ Asian countries like Taiwan, China, Korea and Japan have reported high prevalence rates of 70-80% for nosocomial MRSA. ^2, 3^ In the Philippines, the MRSA rates have steadily increased since 2004 and remained above 50% since 2010, while resistance rates to antibiotics other than beta-lactams are low (Figure 1A-C).

**Figure 1.**
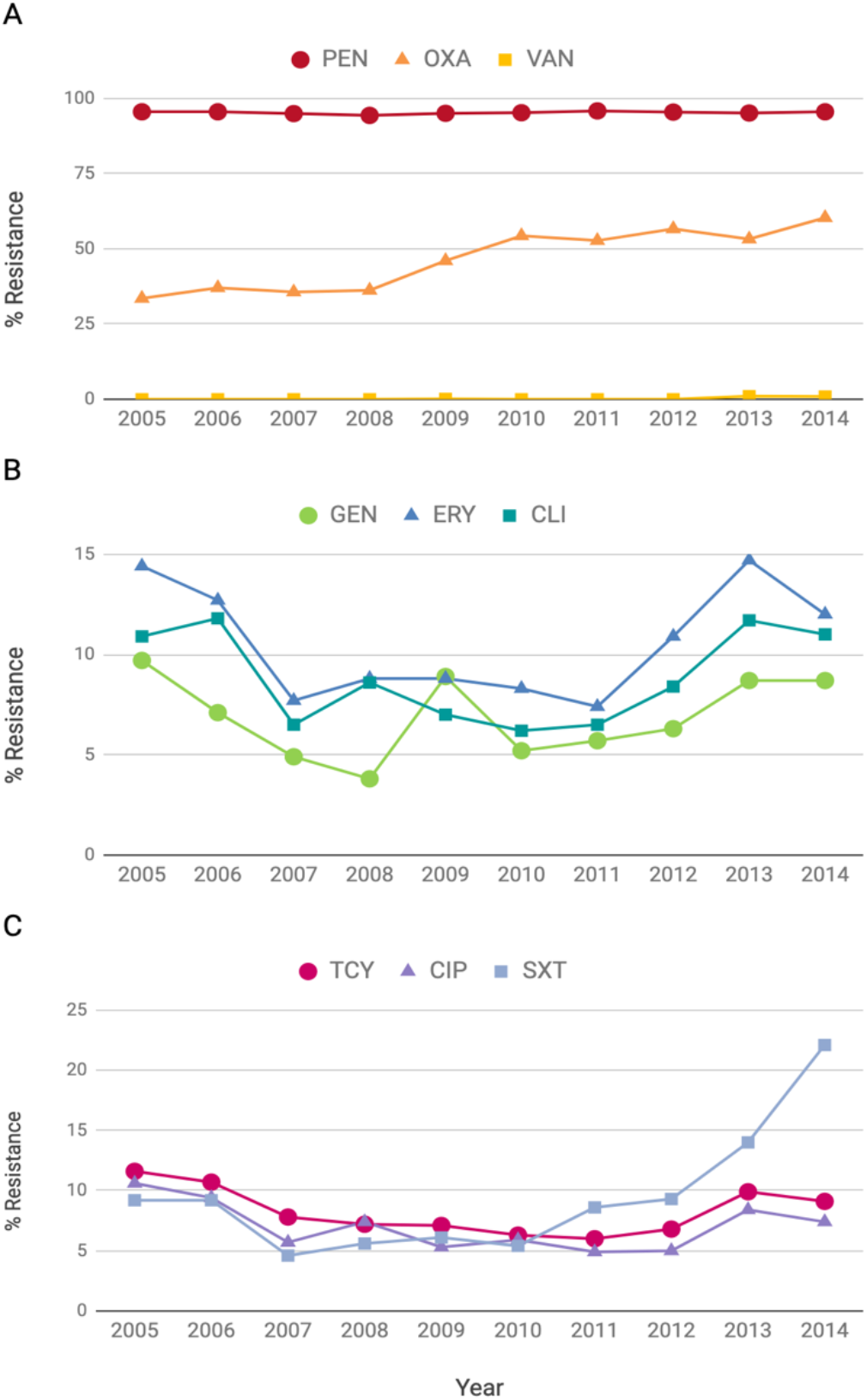
Yearly resistance rates of *S. aureus* isolates 2005-2014 for 9 antibiotics. **A)** PEN: penicillin; OXA: oxacillin; VAN: vancomycin, **B)** GEN: gentamicin; ERY: erythromycin; CLI: clindamycin, **C)** TCY: tetracycline; CIP: ciprofloxacin; SXT: trimethoprim-sulfamethoxazole.

Several notable epidemic clones have spread across Asia, known by their Multi-Locus Sequence Type (MLST) as ST30 (Philippines, Taiwan, Hong Kong, Singapore, Japan, Malaysia and Kuwait), ST239 (Philippines, Thailand, Korea, Vietnam, Taiwan, and India), ST5 (Philippines, Taiwan, Hong Kong, Sri Lanka, Korea and Japan), ST59 (Taiwan, Hong Kong, Vietnam, Sri Lanka and China), and ST72 (Korea). ^3–5^ MRSA strains have emerged independently in the context of these different epidemic clones by acquiring the staphylococcal cassette chromosome *mec* (SCC*mec*) that carries the *mec*A or *mec*C genes, ^6^ which confer resistance to methicillin and most β-lactam antibiotics. Importantly, some MRSA clones have also acquired resistance to vancomycin, the first-line antibiotic treatment for severe MRSA infections in hospitals. ^7^ However, vancomycin resistance has remained very low in the Philippines. ^8^

Current infection control in the Philippines includes cohorting of patients with MRSA infection and laboratory-based surveillance to determine the antimicrobial susceptibility pattern. However, the MRSA burden can be partially attributed to pathogen-specific characteristics of the circulating clones, like antibiotic resistance and virulence genes. ^1^ Hence, a good understanding of the genomic epidemiology of MRSA in the Philippines can aid in the control and management of MRSA infections.

## Methods

### Bacterial Isolates

A total of 6211 *S. aureus* isolates data were collected by the Philippine Department of Health - Antimicrobial Resistance Surveillance Program’s (DOH-ARSP) during the period of January 2013 to December 2014. Isolates found to be resistant to oxacillin (i.e., MRSA) were subsequently referred to the ARSP reference laboratory (ARSRL) for confirmation. Out of the 412 and 384 isolates referred in 2013 and 2014, respectively, a total of 118 MRSA isolates representing 17 sentinel sites were selected for whole-genome sequencing based on their resistance profile (Table 1) with the following criteria: i) referred to ARSRL in 2013-2014; ii) complete resistance profile (i.e., no missing susceptibility data); iii) overall prevalence of the resistance profile in the ARSP data (including both referred and non-referred isolates); iv) geographical representation of different sentinel sites. The number of isolates included from each sentinel site proportional to their relative abundance and estimated from (n/N)*100 (rounded up), where n is the total number of isolates from one site, and N is grand total of isolates; v) when both invasive and non-invasive isolates representing a combination of resistance profile, sentinel site and year of collection were available, invasive isolates (i.e. from blood, or cerebrospinal, joint, pleural and pericardial fluids) were given priority. We utilized a proxy definition for “infection origin” whereby patient first isolates collected in the community or on either of the first two days of hospitalization were categorized as community-acquired infection isolates (CA), while isolates collected on hospital day three or later were categorized as hospital-acquired (HA) infection isolates. ^9^

**Table 1.** Number of *S. aureus* and MRSA isolates analysed by the ARSP and referred to the ARSRL during 2013 and 2014, isolates submitted for whole-genome sequencing, and high-quality MRSA genomes obtained, discriminated by sentinel site and AMR profile.

|  | Number of Isolates |  |  |
| --- | --- | --- | --- |
|  | 2013 | 2014 | Total |
| **Total ARSP** |  |  |  |
| *S. aureus* | 2682 | 3529 | 6211 |
| MRSA | 1421 (53%) | 2128 (60%) | 3549 (57%) |
| **Referred to ARSRL** |  |  |  |
| *S. aureus* | 412 | 384 | 796 |
| MRSA | 381 (92%) | 354 (92%) | 735 (92%) |
| **MRSA submitted for WGS** | 57 | 61 | 118 |
| **MRSA high-quality genomes** | 55 | 61 | 116 |
| *By sentinel site* |  |  |  |
| **BGH** | 6 | 3 | 9 |
| **BRH** | 0 | 1 | 1 |
| **CMC** | 2 | 2 | 4 |
| **CVM** | 6 | 4 | 10 |
| **DMC** | 4 | 3 | 7 |
| **EVR** | 2 | 3 | 5 |
| **FEU** | 3 | 3 | 6 |
| **GMH** | 2 | 3 | 5 |
| **JLM** | 2 | 4 | 6 |
| **MAR** | 4 | 5 | 9 |
| **MMH** | 1 | 2 | 3 |
| **NKI** | 3 | 4 | 7 |
| **NMC** | 3 | 3 | 6 |
| **SLH** | 0 | 1 | 1 |
| **STU** | 5 | 5 | 10 |
| **VSM** | 12 | 13 | 25 |
| **ZMC** | 0 | 2 | 2 |
| *By AMR profile* |  |  |  |
| PEN OXA | 45 | 59 | 104 |
| PEN OXA SXT | 7 | 2 | 9 |
| PEN OXA GEN ERY CLI TCY CIP | 2 | 0 | 2 |
| PEN OXA ERY | 1 | 0 | 1 |

### Antimicrobial Susceptibility Testing (AST)

All *S. aureus* from this study were tested for antimicrobial susceptibility to 14 antibiotics representing 8 different classes: penicillin, oxacillin, cefoxitin, chloramphenicol, sulfamethoxazole/trimethoprim, gentamycin, erythromycin, clindamycin, tetracycline, ciprofloxacin, levofloxacin, rifampin, linezolid, and vancomycin. Antimicrobial susceptibility of the isolates was determined at the ARSRL using Kirby-Bauer disk diffusion method and/or Vitek 2 Compact automated system (BioMerieux). The zone of inhibition and minimum inhibitory concentration of antibiotics were interpreted according to the 26^th^ edition of the Clinical and Laboratory Standard Institute (CLSI) guidelines. ^10^

### DNA Extraction and Whole-Genome Sequencing

A total of 118 MRSA isolates were shipped to the Wellcome Trust Sanger Institute for whole-genome sequencing. DNA was extracted from a single colony of each isolate with the QIAamp 96 DNA QIAcube HT kit and a QIAcube HT (Qiagen; Hilden, Germany). DNA extracts were multiplexed and sequenced on the Illumina HiSeq platform (Illumina, CA, USA) with 100-bp paired-end reads. Raw sequence data were deposited in the European Nucleotide Archive (ENA) under the study accession PRJEB17615. Run accessions are provided on the Microreact projects.

### Bioinformatics analysis

Genome quality was evaluated based on metrics generated from assemblies, annotation files, and the alignment of the isolates to the reference genome of *Staphylococcus aureus* subsp *aureus* strain TW20 (accession FN433596), as previously described. ^11^ Annotated assemblies were produced as previously described. ^12^ A total of 116 high-quality *S. aureus* genomes were included in this study.

We derived *in silico* the multi-locus sequence type (MLST), the *spa* type and SCC*mec* type of the isolates from the whole genome sequences. The sequence types (ST) were determined from assemblies with Pathogenwatch (https://pathogen.watch/) or from sequence reads with ARIBA ^13^ and the *S. aureus* database hosted at PubMLST ^14^. The *spa* type was inferred with spaTyper v1.0 ^15^. The SCC*mec* type was derived from sequence reads with SRST2 ^16^ and the database available at http://www.staphylococcus.net/Pages/indexEN.html.

Evolutionary relationships between isolates were inferred from single nucleotide polymorphisms (SNPs) by mapping the paired-end reads to the reference genomes of *S. aureus* strains TW20 (FN433596) or ILRI_Eymole1/1 (NZ_LN626917), as described in detail previously ^11^. Mobile genetic elements (MGEs) were masked from the alignment of pseudogenomes. For the CC30 phylogeny recombination regions detected with Gubbins ^17^ were also removed. SNPs were extracted with snp_sites ^18^ and maximum likelihood phylogenetic trees were generated with RAxML ^19^ based on the generalised time reversible (GTR) model with GAMMA method of correction for among-site rate variation and 100 bootstrap replications. The tree of 7821 global *S. aureus* genomes available at the European Nucleotide Archive with geolocation and isolation date was inferred using an approximately-maximum likelihood phylogenetic method with FastTree. ^20^

Known AMR determinants and the Panton-Valentine leukocidin (PVL) *lukF-PV* and *lukS-PV* genes were identified from raw sequence reads using ARIBA ^13^ and a curated database of known resistance genes and mutations. ^21^ Genomic predictions of resistance were derived from the presence of known antimicrobial resistance genes and mutations identified in the genome sequences. The genomic predictions of AMR (test) were compared to the phenotypic results (reference) and the concordance between the two methods was computed for each of 8 antibiotics (928 total comparisons). Isolates with either a resistant or an intermediate phenotype were considered resistant for comparison purposes.

*S. aureus* assemblies were also analysed using Pathogenwatch (https://pathogen.watch/), which infers trees based on genetic similarity, and predicts genotypic AMR. All project data, including inferred phylogenies, AMR predictions and metadata were made available through the web application Microreact (http://microreact.org).

## Results

### Demographic and Characteristics of the MRSA Isolates

Of the 118 MRSA isolates submitted for WGS, 116 isolates were confirmed as *S. aureus in silico*, while 2 isolates were identified as *Staphylococcus argenteus*, and were not included in the downstream analyses. The age range of the patients was <1 to 90 years old, with 20% (N=23) of the isolates from patients from the age group of <1 year old (Table 2). Among the 116 isolates, 56% were recovered from male patients (N=66) and 44% from females (N=50). Since invasive isolates were prioritized, the most common specimen source was blood (62%, N=72), followed by wound (19%, N=22). The majority of the infections (68%, N=79) were classified as community-associated MRSA.

**Table 2.** Demographic and clinical characteristics of 116 MRSA isolates. Invasive isolates were considered as those obtained from specimen types marked with an asterisk (*).

| **Characteristic** | **No. Isolates** |
| --- | --- |
| **Sex** |  |
| Male | 66 |
| Female | 50 |
| **Age (in years)** |  |
| <1 | 23 |
| 1-4 | 9 |
| 5-14 | 16 |
| 15-24 | 14 |
| 25-34 | 8 |
| 35-44 | 17 |
| 45-54 | 14 |
| 55-64 | 9 |
| 65-80 | 1 |
| >=81 | 5 |
| **Patient Type** |  |
| in | 104 |
| out | 12 |
| **Specimen Origin** |  |
| Community-acquired | 79 |
| Hospital-acquired | 37 |
| **Specimen Type** |  |
| Abdominal Fluid* | 1 |
| Abscess | 4 |
| Aspirate | 2 |
| Blood* | 72 |
| Bone | 1 |
| Cerebrospinal Fluid* | 2 |
| Pericardial Fluid* | 1 |
| Tracheal Aspirate | 5 |
| Urine | 2 |
| Wound | 22 |
| Others | 1 |

### Concordance Between Phenotypic and Genotypic AMR

Isolates were tested for susceptibility to 14 antibiotics representing 8 different classes. All isolates were susceptible to vancomycin and linezolid and resistant to penicillin, oxacillin and cefoxitin, consistent with the presence of the *blaZ* and *mecA* genes (Table 3). Nine isolates were resistant to cotrimoxazole, which was associated with the presence of the *dfrG* gene. Two isolates were multi-drug resistant (MDR) and carried several genes and mutations, with resistance to penicillin (*blaZ, mecA*), oxacillin (*mecA*), cefoxitin (*mecA*), gentamicin (*aacA-aphD*), erythromycin (*ermC, msrA*), clindamycin (*ermC*), tetracycline (*tetM*, *tetK*), ciprofloxacin and levofloxacin (*GyrA*_S84L, *GyrA*_G106D, and *GrlA*_S80F mutations), chloramphenicol (*catA1*) and rifampicin (*rpoB*_H481N). The *IleS* gene that confers resistance to mupirocin, and the *sdrM* gene conferring resistance to norfloxacin were identified in 3 and 23 isolates, respectively. However, mupirocin was not tested in the laboratory, and norfloxacin was only tested against isolates from urine specimens. Hence, these two antibiotics were not included in the concordance analysis.

**Table 3.**
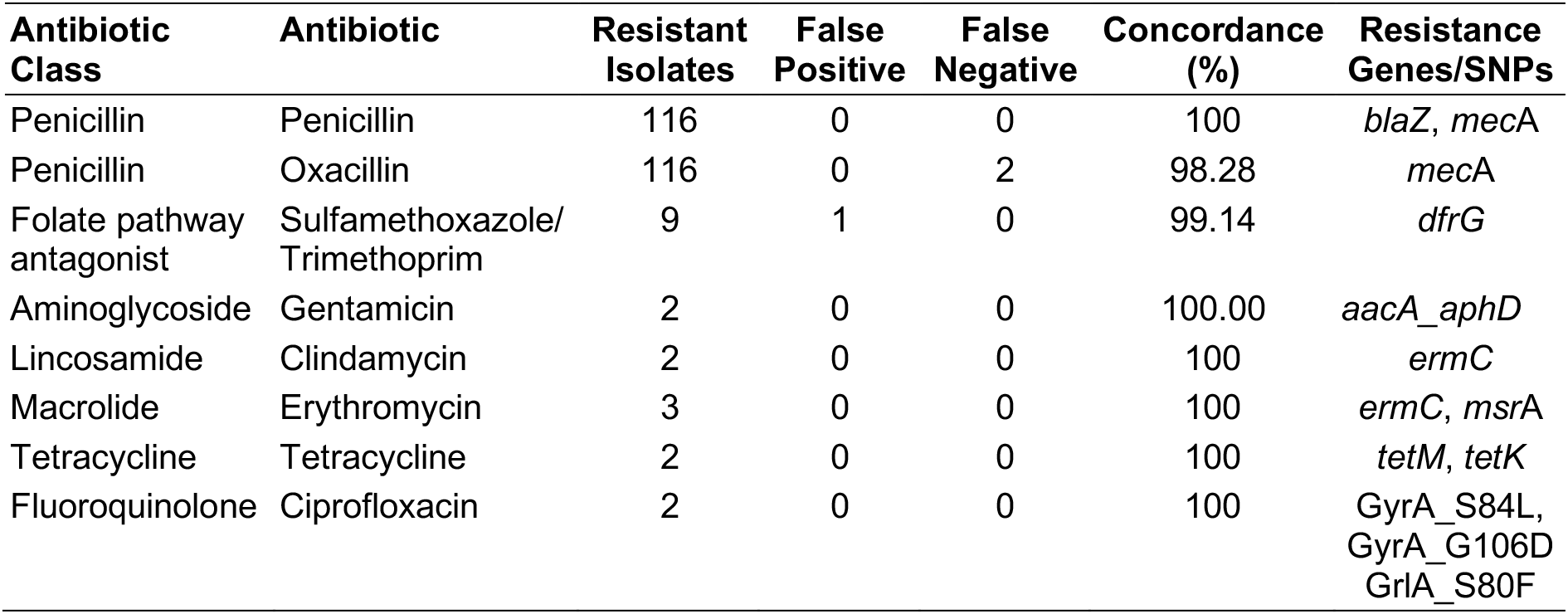
Comparison of genomic predictions of antibiotic resistance with laboratory susceptibility testing at the ARSRL.

Comparisons between phenotypic and genotypic data are presented for 8 key antibiotics in 7 classes (Table 3). The overall concordance for the 928 comparisons was 99.68%, and the concordance for individual antibiotics was above 98% in all cases (Table 3). The notable exceptions were two false negative calls for oxacillin resistance, i.e. isolates confirmed to be phenotypically resistant but without the *mec*A resistance gene. Conversely, one isolate was falsely predicted to be resistant to sulfamethoxazole/trimethoprim based on the presence of the *dfrG* gene.

## Genotypic findings

### *In silico* genotyping

Multi-locus sequence type (MLST), *spa* type, and SCC*mec* type were predicted *in silico* from the whole-genome sequence data of the 116 MRSA isolates. A total of 18 sequence types (ST) were identified, with 74.1% (N=86) of the isolates belonging to clonal complex 30 (CC30), distributed between ST30 (N=81), its single-locus variant ST1456 (N=2), and three ST30 genomes showing novel *aroE* (N=2) and *yqiL* (N=1) alleles. Clonal complex 5 (CC5) was represented by nine genomes, followed by ST834 (N=6). The most prevalent of the 29 different *spa* types identified was t019 (62%), which coincided with genomes assigned to CC30. The *spa* types identified for the CC5 genomes, were t002 (N=5), t105 (N=3) and t067 (N=1). The vast majority of the SCC*mec* cassettes identified in our genomes belonged to type-IV (N=108, 93.1%), followed by type-III (N=2, 1.7%). The SCC*mec* type could not be determined for four genomes. Details of the number and most common ST and *spa* types found in each of the sentinel sites are shown on Table 4. Altogether, the typing results derived from the genome sequences showed that CC30-*spa*-t019-SCCmec-IV was the most prevalent MRSA clone in our retrospective collection (N=67, 57.8%).

**Table 4.**
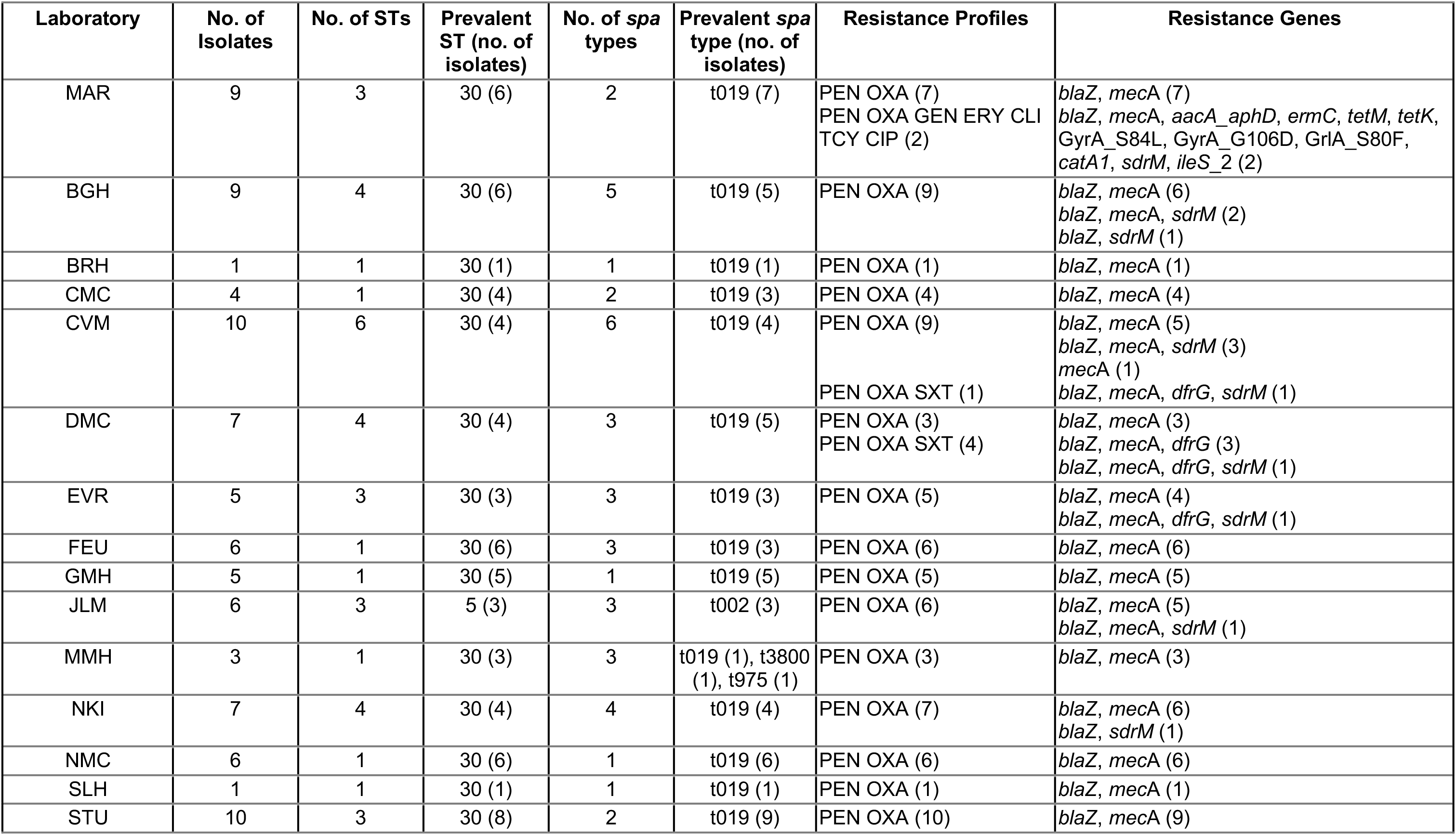
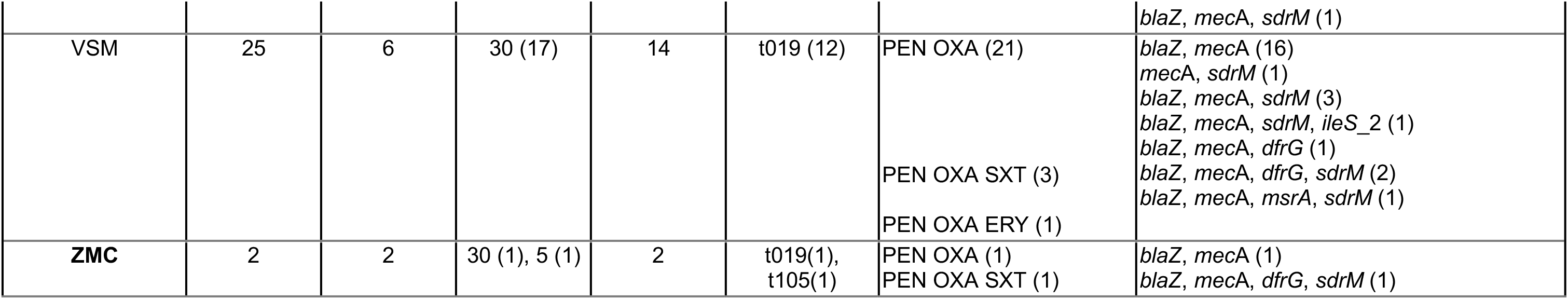
Distribution of isolates, sequence types (STs), *spa* types, resistance profiles and AMR genes and mutations across the 17 sentinel sites.

### Population structure of MRSA in the Philippines

The phylogenetic tree shows that the population was composed of discrete clades that matched the ST distribution and that were separated by long branches (Figure 2A), in agreement with the clonal population previously described for *S. aureus*. ^22^ The largest clade represented by CC30-*spa*-t019-SCC*mec*-IV was characterized by a broad geographical distribution across the 17 sentinel sites in this dataset (Table 4). It was found among both community-associated (CA), and healthcare-associated (HA) isolates obtained from at least 11 different specimen types. WGS revealed distinct major sublineages within the CC30 clade (Figure 2B), none of which displayed strong phylogeographic signal. Both genes *lukS-PV* and *lukF-PV* encoding the Panton-Valentine leukocidin (PVL) were found in 75 of the 86 CC30 genomes (Figure 2B), indicating that the majority (87.2%) are PVL-positive (Figure 2B).

**Figure 2.**
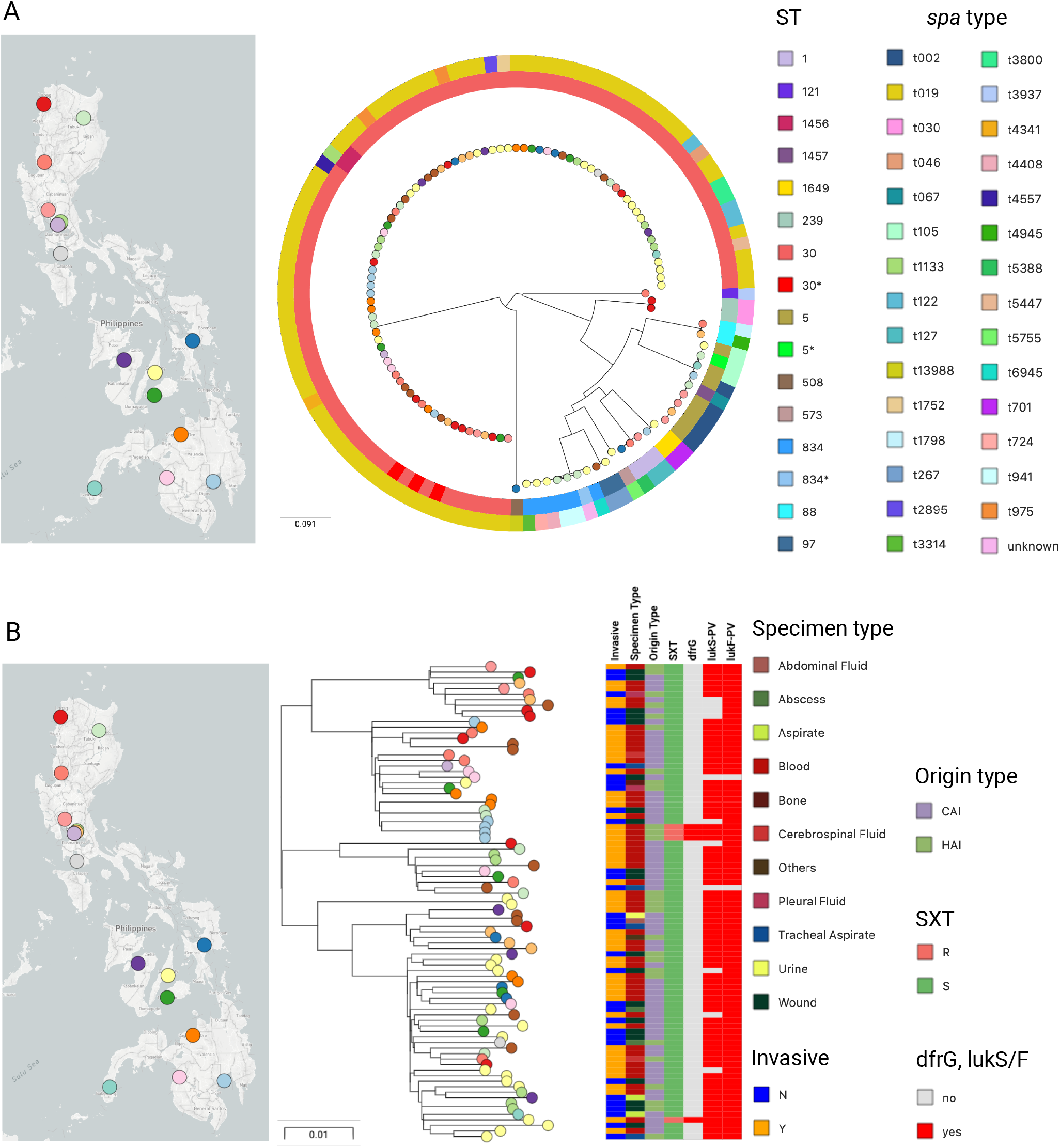
Genomic surveillance of *S. aureus* from the Philippines 2013-2014. **A)** Phylogenetic tree of 116 MRSA isolates from the Philippines, inferred with RAxML from 96,514 SNPs sites obtained after mapping the genomes to the complete genome of strain TW20 (ST239) and masking regions with mobile genetic elements. The tree leaves are coloured by sentinel site. Inner ring: Sequence Type. Outer ring: *spa* type. The data is available at https://microreact.org/project/ARSP_SAU_2013-2014. **B)** Phylogenetic tree of 86 CC30 isolates inferred with RAxML from 4,780 core SNPs obtained after mapping the genomes to the complete genome of strain Eymole1 (ST30) and masking mobile genetic elements and recombination regions from the alignment. The data is available at https://microreact.org/project/ARSP_SAU_CC30_2013-2014. The scale bars represent the number of single nucleotide polymorphisms (SNPs) per variable site.

Two additional epidemic clones were identified, CC5 and ST239 (CC8). The nine CC5-SCC*mec*-typeIV were mostly from pediatric patients (N=6, 71% compared to 20% of the entire dataset), and clustered largely according to their *spa* type, but displayed no clear phylogeographic distribution. Two isolates from different, distant locations carried both *lukS-PV* and *lukF-PV* genes (PVL-positive). The two ST239 isolates were from the same patient, *spa* type t030, SCC*mec*-typeIII, PVL-negative, and multi-drug resistant.

The nine isolates with resistance to sulfamethoxazole/trimethoprim belonged to four different locations (CVM, DMC, VSM, and ZMC), and four different clones (CC30, CC5, ST1649, and ST834), which suggests that resistance to this antibiotic has emerged independently (Table 4). The isolates were obtained from blood (N=8) and abscess (N=1) and, interestingly, mostly from pediatric patients (6 out of 7 patients, 85.7% compared to 46.6% pediatric patients in the total dataset).

### MRSA from the Philippines in global context

We placed the genomes from our retrospective collection in the global context of 7821 contemporary *S. aureus* public genomes available from sequence data archives with linked geographic and temporal information, and collected between 2010 and 2017. This collection of public genomes represents 57 countries and 379 STs, but it is heavily biased towards genomes from Europe (N=3556) and USA (N=3241) and belonging to the epidemic clones ST8 (N=2343), ST22 (N=1526) and ST5 (N=720) prevalent in those regions (Figure 3A). CC30-*spa-*t019-SCC*mec*-IV-PVL+ MRSA genomes from the Philippines formed a discrete cluster within ST30 together with a small number of genomes from the USA (N=5), the UK (N=3), and Germany (N=1, Figure 3B).

**Figure 3.**
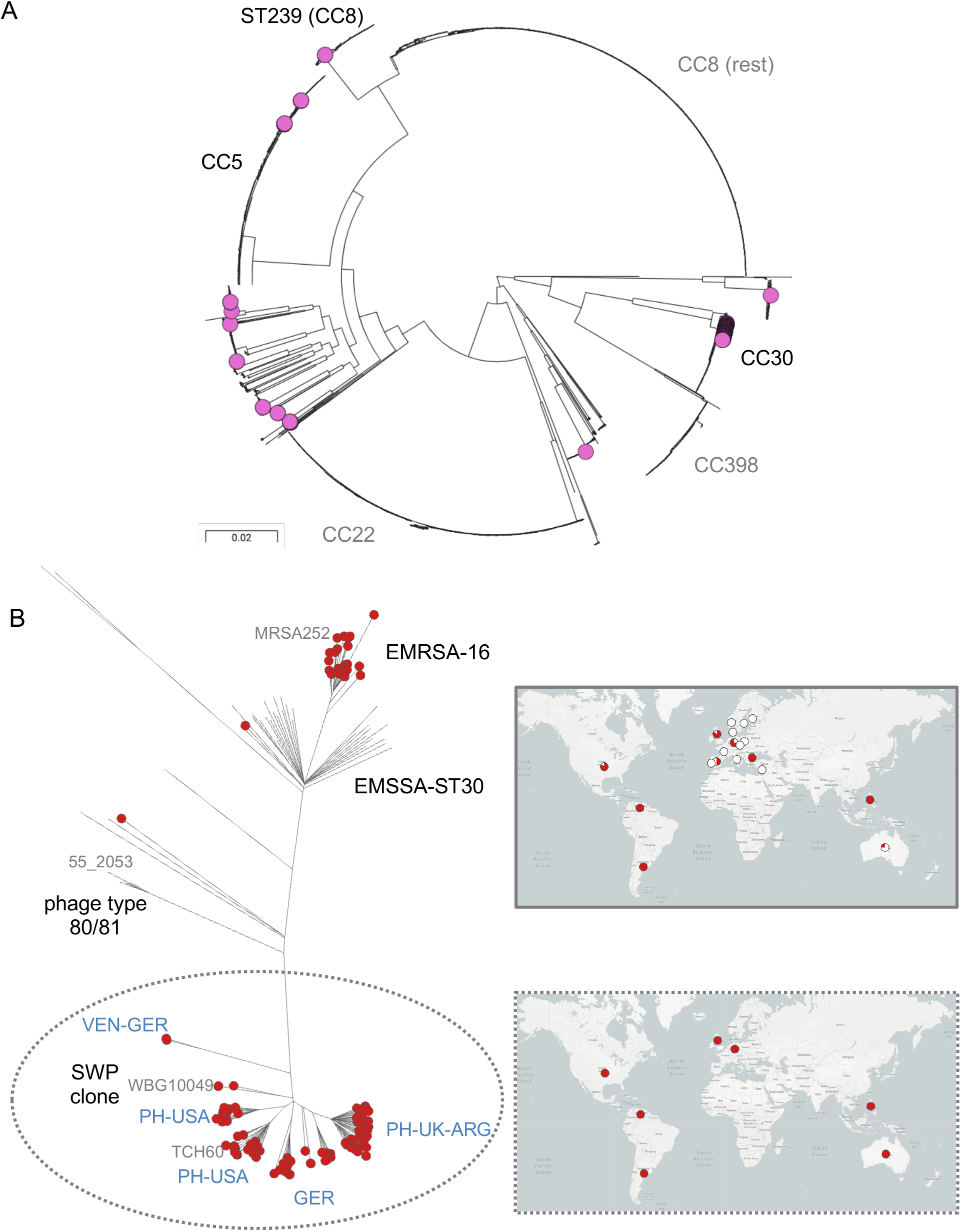
*S. aureus* from the Philippines in global context. **A)** Phylogenetic tree of 7821 *S. aureus* isolates from the Philippines (N=116, this study) and from 57 other countries inferred with FastTree from 485,031 SNPs positions obtained after mapping the genomes to the complete genome of strain TW20 and masking regions with mobile genetic elements. The magenta tree nodes indicate the genomes from this study. The major lineages (CCs and STs) are labelled in black if represented by genomes of this study, or in grey if they are not. The scale bars represent the number of single nucleotide polymorphisms (SNPs) per variable site. The data is available at https://microreact.org/project/Global_SAU. **B)** Pathogenwatch tree of 166 CC30 genomes comprising 86 genomes from this study and 80 available global genomes. Red nodes: MRSA, otherwise MSSA. Top map: Geographical distribution of 166 genomes. Bottom map: geographical distribution of the genomes within the dotted line. The tree is annotated with previously described CC30 clones (black) and their representative reference genomes (grey). The collection is available at https://pathogen.watch/collection/vm2mfuj83zbx-arsp-sau-cc30-2013-2014-global-context.

A number of successful pandemic clones have emerged within CC30, such as the methicillin-sensitive (MSSA) phage type 80/81, ^23^ the MRSA South-West Pacific clone (SWP, ^24^), the hospital-endemic EMRSA-16 (ST36, ^25^), and EMSSA-ST30. ^26^ We contextualized the Philippine MRSA genomes from this study with available genomes from these clones ^25–29^ using Pathogenwatch, which showed that the Philippine genomes form a lineages related to, but distinct from the SWP clone, representing a new diversification from this clone (Figure 3B). In addition, the genomes from the Philippines clustered with genomes from USA, Germany, the UK and Argentina, indicating that this epidemic diversification from the SWP clone has been accompanied by global dissemination.

## Discussion

In this study we combined WGS and laboratory-based surveillance to characterize MRSA circulating in the Philippines in 2013 and 2014. High levels of concordance between phenotypic and genotypic resistance was observed for all antibiotics tested (Table 3). This has been previously reported for *S. aureus* collections from the UK and Europe, ^26, 30^ but our results, the first from the Philippines, shows that there are no significant gaps in the epidemiology of known resistance mechanisms in this country. The integration of laboratory and WGS data showed the independent acquisition of resistance to sufamethoxazole/trimethoprim mostly in pediatric patients with invasive infections. This is likely due to the selective pressure of antibiotic use, as co-trimoxazole was historically one of the first line antibiotics given to pediatric patients with pneumonia as recommended by the Department of Health in the Philippines in the 1990s and it is currently the recommended first line antibiotic for skin and soft tissue MRSA infections in pediatric patients by the Philippines National Antibiotic Guideline for 2017.

Few studies described the molecular epidemiology of MRSA in the Philippines. We found that the MRSA population captured in 2013 and 2014 was composed of a limited number of genetic lineages, and dominated by CC30-spa-t019-SCC*mec*-IV-PVL+ (Figure 2A). Community-acquired CC30-*spa*-t019-PVL+ MRSA was previously reported for one hospital in the Philippines and the period between 2004-2006, and also from other countries in South East Asia, with potential for clonal expansion. ^3, 4^ This suggests that the increase in the burden MRSA observed in the Philippines since 2004 (Figure 1A) may be linked to the expansion of this clone (Figure 2A). Extending the period of the retrospective sequencing survey would provide an in-depth insight into the dynamics of this clone, but was beyond the scope of this study. The lack of a clear phylogeographic structure in this established clone (Figure 2B) may be related to its ability to disseminate both within hospitals and into the community, and within the community followed by nosocomial transmission, ^31^ coupled with the dense population of the Philippines. However, the ARSP hospital-based surveillance would need to be complemented with community surveillance to understand the dynamics of MRSA in the population at large. In addition, only a small number of public global genomes were found to be closely related to the CC30 genomes in this study (Figures 3A and B), highlighting the paucity of WGS data from continents other than North America and Europe, which would enhance our understanding of the global epidemiology of this clone.

The genotypic characterization of circulating MRSA strains, together with the phenotypic and epidemiological data, led to the identification of several global epidemic clones and revealed the lack of strong phylogeographic structure in the population from patients admitted to healthcare facilities in a country with high burden of MRSA. This supports the implementation of interventions aimed to reducing the burden of disease in the general population. ^4^ Targeted eradication interventions may be useful for individual hospitals where high-risk epidemic clones such as ST239 may cause local outbreaks. Our results represent the first comprehensive genomic survey of *S. aureus* in the Philippines, which bridges the gap in genomic data from the Western Pacific region and will constitute the genetic background to contextualize ongoing prospective surveillance to inform infection control.

## Acknowledgements

Funding provided by Newton Fund, Medical Research Council (UK), Philippine Council for Health Research and Development.

